# Careers in Scientific Editing: Results from the 2024 Scientific Editors Network (ScENe) Survey

**DOI:** 10.64898/2026.10.05.756087

**Authors:** Heather Aldrich, Jennifer Barr, Becky James, Robert Lawrence, Megan Mayerle, Ann Sutton

## Abstract

Funding and publishing scientific research has become increasingly competitive, highlighting a growing need for clear and accurate scientific communication. Scientific editors are professionals who help ensure that grant proposals and research manuscripts meet the standards of funding agencies and peer-reviewed journals while effectively communicating research significance to diverse audiences. However, detailed information characterizing the scientific editing profession is scarce. While some organizations have conducted salary surveys, to our knowledge, no comprehensive survey of the scientific editing field has been conducted. Here, we report findings from a 2024 comprehensive survey of 132 scientific editors. We collected data on demographics and education, career experience and current employment, skills and types of editing, and outcomes and satisfaction. Our data reveal that scientific editors have homogenous demographic characteristics but considerable variability in job titles, employment mechanisms, and career pathways. As the responsibilities of many scientific editors, including supporting proposal development, applying best practices in grantsmanship, and identifying strategies to enhance competitiveness for research funding, overlap with those of research development professionals, our findings provide considerations for further incorporating or defining the role of scientific editors within research development offices. Our results also suggest avenues for scientific editors to start or advance their careers. We anticipate that our results will serve as a foundation for discussions on the future directions of the scientific editing profession as well as areas for future research and professional development.

## Introduction

The ability to convey scientific data and ideas in a clear and accessible way is an essential skill for scientists, as it improves communication and fosters collaboration within and between scientific fields^1–3^ and improves the ability of specialists and non-specialists to understand the implications of the research.^4^ Indeed, well-written publications are more likely to be cited^4–6^ and clearly written grant proposals are more likely to be funded.^7,8^ In contrast, poorly written studies can lead to misunderstandings or even misrepresentation of research findings by scientists, as well as the general public.^9–11^ Amid intensifying competition for research funding and for publication in high-impact journals, the need for clear, compelling, and accurate scientific communication has never been greater.

Many scientists rely on scientific editors to support and improve their writing.^12–14^ A scientific editor’s primary role is to conduct, or mentor others in, substantive editing of research grant proposals and/or manuscripts published in peer-reviewed journals. Substantive editing goes beyond copyediting to address issues of organization, design, and style, ensuring that documents are not only technically correct but also consistent, logically structured, and compelling.^15^ Scientific editors must be adept at working within tight deadlines and balancing the needs of multiple stakeholders, including researchers, funding agencies, and publishers. Scientific editors also play an essential role in ensuring that research proposals and manuscripts meet the exacting standards of funding agencies and peer-reviewed journals while effectively communicating the significance, innovation, and rigor of research in an engaging way to diverse audiences. When working on grants, scientific editors tend to focus their efforts on developing proposals, offering grantsmanship support, and contributing to a strategy that enhances competitiveness. Unlike research development professionals, scientific editors are often not responsible for disseminating information about funding opportunities, making decisions regarding internal funding, providing strategic guidance at the institutional level, or facilitating collaborations between investigators. Thus, although there are many similarities in job function between scientific editors and research development professionals, the specific emphasis on proposal development by scientific editors demonstrates that they are an important subset of the research development profession.

Despite the importance of scientific editing to grant funding and publishing, there are limited empirical data characterizing the field and its workforce as a distinct profession. Some professional organizations related to scientific editing have conducted salary surveys, such as the American Medical Writers Association (AMWA)^16^ and the National Association of Research Development Professionals (NORDP).^17^ However, to our knowledge, no comprehensive survey focused on scientific editors has been conducted. Here, we report findings from a 2024 survey of individuals whose jobs include scientific editing. Our data provide a benchmark for the field of scientific editing and demonstrate the lack of a career path, training, and support for those interested in entering the field. By synthesizing the findings of our survey and related literature, this article aims to provide a comprehensive overview of the state of the scientific editing profession, highlight its importance to the research enterprise and the field of research development, and identify areas for future research and professional development. Our findings may provide important considerations for the strategic expansion or growth of research development offices to incorporate training and career paths for individuals who perform scientific editing.

## Methods

### Survey development

The Scientific Editors Network (ScENe) is a professional organization with the mission, “to provide an inclusive, welcoming forum for scientific editors to share skills, knowledge, and resources; discuss strategies to overcome challenges; and improve science communication.”^18^ The 2024 ScENe survey was developed by six members of ScENe who live across the United States (U.S.) and have a variety of titles and experience within the field of scientific editing. The survey was then pilot tested with four other ScENe members who provided additional perspectives on scientific editing roles and work environments. Following survey development best practices,^19,20^ face validity of the survey was evaluated by the author team who assessed the questionnaire for clarity, style, ease of understanding, and layout. The additional content experts who pilot tested the survey reviewed the tool for content validity including readability, clarity, comprehensiveness, and agreement on questions to be retained in the final questionnaire. The final survey was created in Qualtrics XM and included 48 quantitative and qualitative questions, with sections covering demographics and education, career experience and current employment, skills and types of editing, and outcomes and satisfaction. The estimated time for completion was approximately 15-20 minutes.

### Participant recruitment

The survey was distributed via a web-based link to email/listserv rosters of seven organizations with members who have roles within scientific writing and editing fields (Table 1). The introductory text encouraged survey recipients to share the survey with colleagues. Reminder emails were sent to each of the email lists approximately two weeks after the initial email. Responses were collected from April 1, 2024, to May 9, 2024.

**Table 1.**
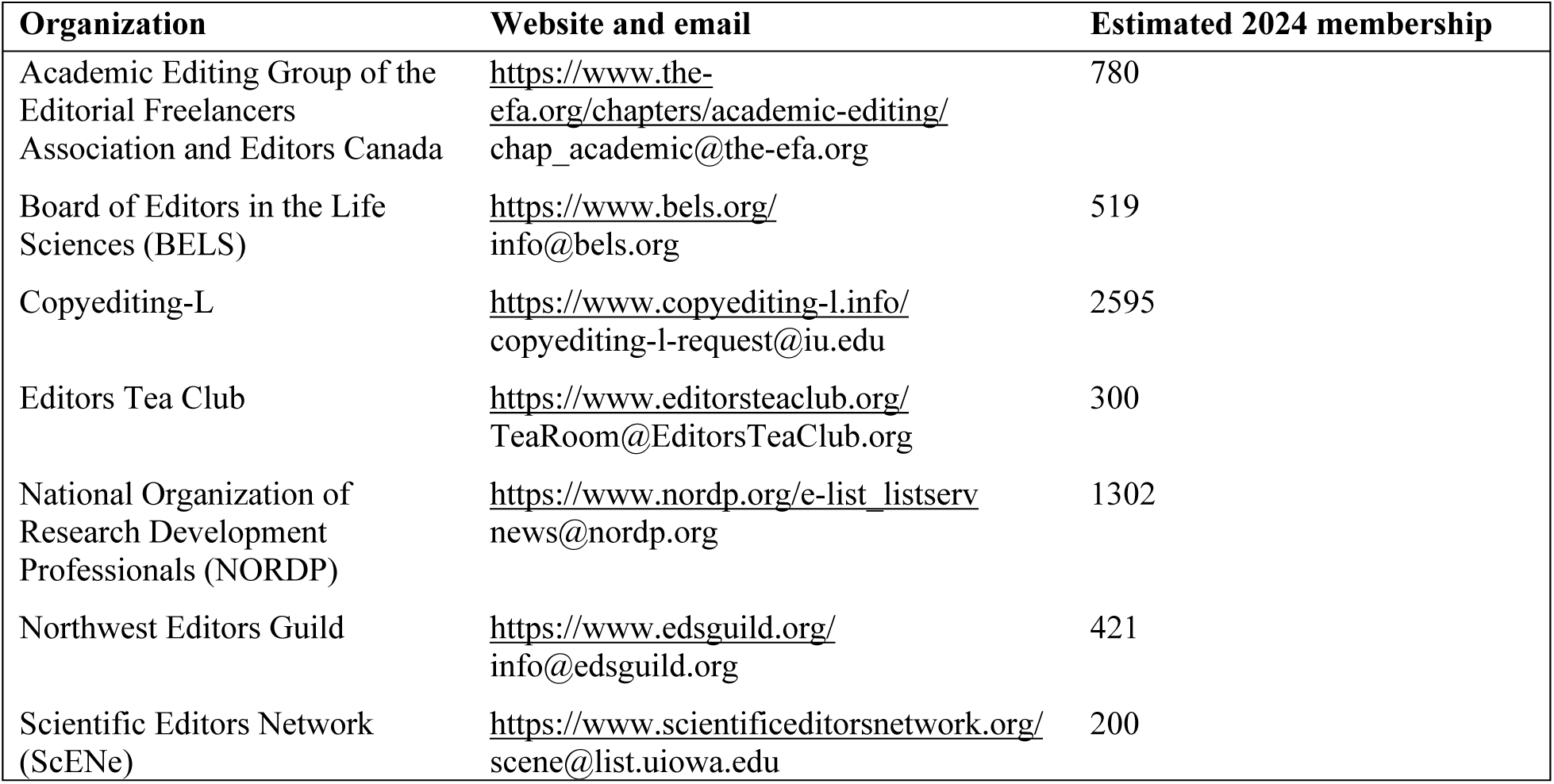
Professional Organizations that Participated in Distributing the Survey.

The inclusion criteria for participating in the study were 1) adults (aged 18 years and over) whose jobs included scientific editing and 2) the ability to complete the survey in English. The first question on the survey was used to screen participants: *Which of the following activities do you perform as part of your job? Please select all that apply: A) Scientific editing of research grants, manuscripts, or other scholarly works; B) Researching grant-writing strategies (e.g., grantsmanship) for scientific/research purposes; C) Providing instruction in scientific writing (e.g., teaching, seminars, mentorship).* Participants who selected one or more of these activities continued to the full survey. Participants were further excluded if they answered no subsequent questions or only responded “Prefer not to answer” to all questions.

The Institutional Review Board at Colorado State University approved the study as exempt human subjects research. Participants did not receive an incentive for completing the survey.

### Data analysis

Upon closure of the survey, results were downloaded to Excel. Descriptive statistics were calculated using Excel for demographic characteristics and questions with categorical responses. Inductive coding was used to qualitatively analyze data from open-ended survey questions and to identify themes and trends in survey responses. Open-ended survey responses were analyzed using the six phases of thematic content analysis.^21,22^ The first three phases of the analysis (becoming familiar with the data, generating initial codes, and sorting data into themes) were done by one author; each author was responsible for analyzing a subset of the survey questions. All authors then discussed and approved the final three phases (reviewing themes, refining themes, and writing up the results).

*Other* responses that consisted of open-ended write-in text were assessed to identify duplicated or missing values from the original survey options. These responses were adjusted to reflect accurate final counts as necessary. All remaining *Other* write-in responses were classified into subcategories. In a few cases, identifying information was removed from responses to preserve participant anonymity, e.g., *Head of Research Development at University NAME* to *Head of Research Development at University*.

## Results

### Survey participants

A total of 132 participants (*N*) met the inclusion criteria and answered at least one question in the survey (Figure 1), with some variation in the number of participants who responded to each survey question (*n*). Of the participants, 111 (84%) provided a response to all questions. Most participants spent approximately 20 minutes on the survey. Here, we provide a summary of the most salient results of the survey. The full survey data, including stated questions, categorized responses, and number of participants (*n*), is provided in Supplemental Table S1.

**Figure 1.**
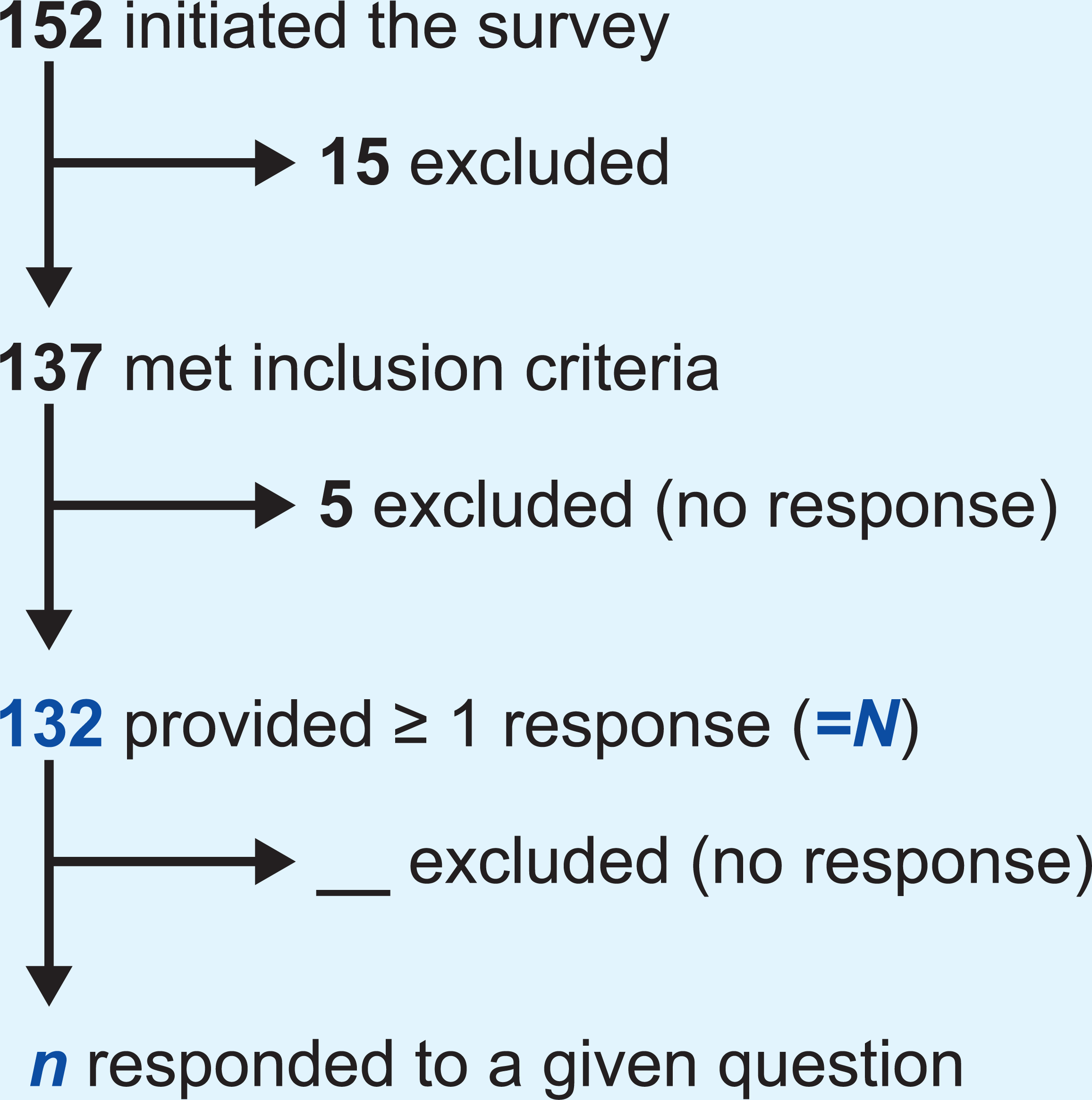
Number of total survey participants (N) and participants to individual survey questions (n), based on inclusion criteria.

### Demographics and education

The demographic characteristics of survey participants are listed in Table 2. The majority of participants were between the ages of 35 to 64 years (80%, combined categories), with the age range of 45-54 years garnering the highest number of responses (30%). Participants overwhelmingly identified as female (83%), and White was the most commonly selected race or ethnicity (87%), followed by Asian (11%).

**Table 2.** Demographic Characteristics of Survey Participants. Bar charts are normalized to the largest value among each category.

| Characteristic | Variables | Percent of participants |
| --- | --- | --- |
| Age, years | 18-24 | 0% |
|  | 25-34 | 9% |
|  | 35-44 | 23% |
|  | 45-54 | 30% |
|  | 55-64 | 27% |
|  | 65+ | 10% |
| Gender | Female | 83% |
|  | Male | 17% |
|  | Prefer not to answer | 1% |
| Race and ethnicity<br>(select all that apply) | American Indian or Alaskan Native | 0% |
|  | Asian | 11% |
|  | Black or African American | 1% |
|  | Hispanic or Latino | 3% |
|  | Native Hawaiian or Pacific Islander | 0% |
|  | White | 87% |
|  | Other | 2% |
|  | Prefer not to answer | 1% |
| Primary residence | US - Northeast | 9% |
|  | US - Atlantic | 18% |
|  | US - Southeast | 17% |
|  | US - Great Lakes | 11% |
|  | US - Midwest/Mountain | 12% |
|  | US - Southwest | 14% |
|  | US - Pacific | 11% |
|  | Canada | 3% |
|  | Other | 5% |
| Education<br>(select all that apply) | High school | 3% |
|  | Associates degree | 2% |
|  | BA/BS | 26% |
|  | MA/MS | 26% |
|  | MPH | 2% |
|  | PhD | 43% |
|  | MD/DO | 0% |
|  | MBA | 1% |
|  | Other | 6% |

Of the 132 survey participants, 92% lived within the U.S., with non-U.S. participants living in Canada, Australia, India, United Kingdom, and Germany. Participants within the U.S. represented all regions (9% Northeast, 18% Atlantic, 17% Southeast, 11% Great Lakes, 12% Midwest/Mountain, 14% Southwest, 11% Pacific).

Participants were asked about the highest level of education completed. PhD was the most common response (43%), followed by bachelor’s (26%) or master’s degree (26%). Participants’ educational background was primarily within the sciences, and included biomedical science (39%), natural sciences (27%), public health (5%), and medicine (2%). Writing and literary disciplines were also well-represented and included English (21%), communications/journalism (17%), and library/information sciences (2%). Notably, 27% of participants selected *Other* and listed fields of study including law, mathematics, social sciences, and education. The survey question allowed for multiple answers, and 10% of participants reported two or more degree areas, usually science paired with writing or literature.

### Current employment

The survey asked participants to provide their formal job titles, prompting a plethora of different responses (Figure 2). Despite the lack of similarity in precise titles, we were able to identify common descriptors, including *editor*, *scientific*, *freelance*, *writer*, *senior*, *medical*, and *research*. The preponderance of participants (80%) indicated employment by a public or private university or research institute, and the balance (20%) identified as exclusively freelancers.

**Figure 2.**
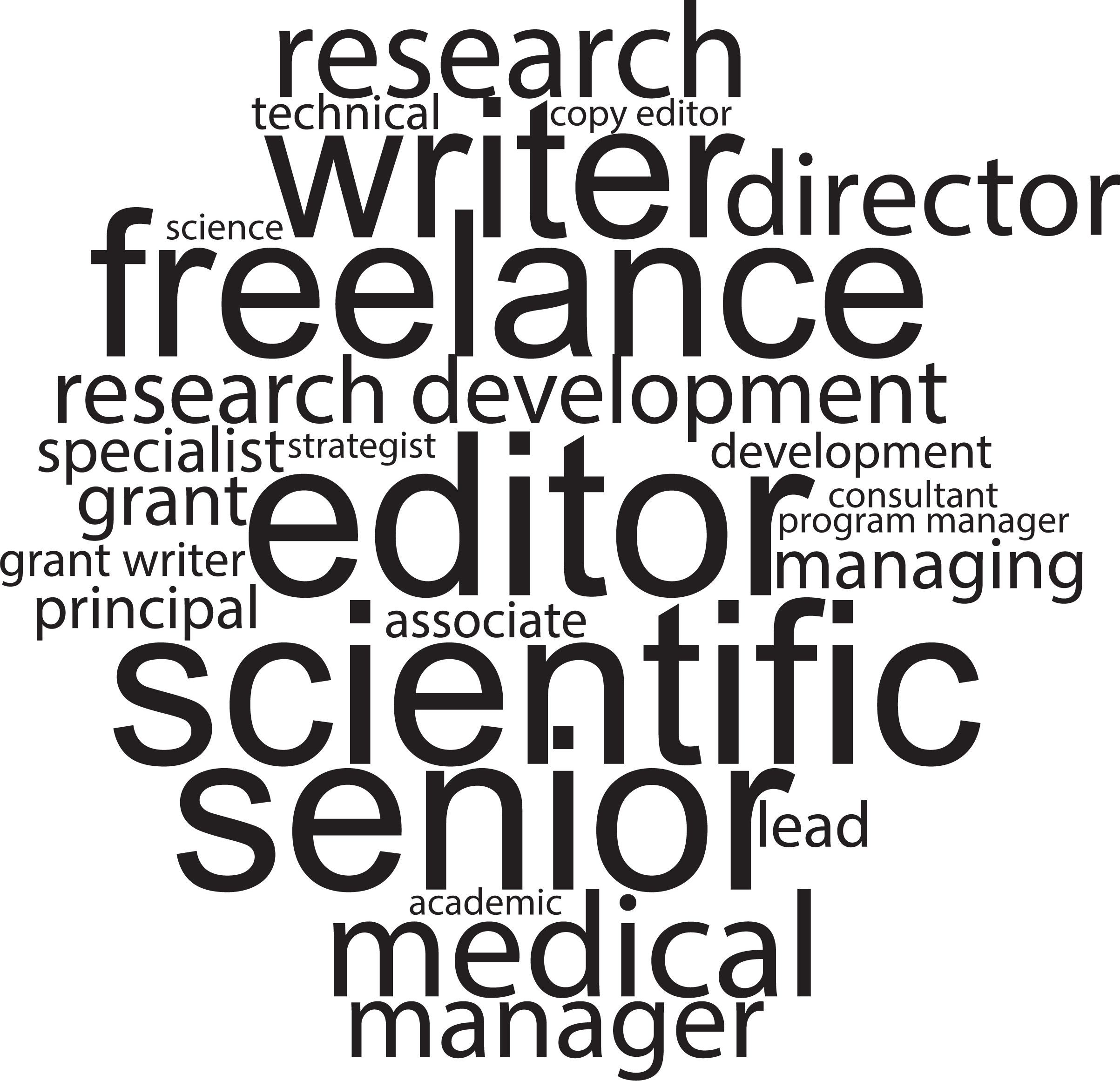
Job title word cloud. The cloud includes words that appeared at least twice in survey participants’ reported job titles.

Within the larger group, the majority (70%) reported being employed full-time. Most (64%) reported working remotely, while nearly 30% had a hybrid arrangement and nearly 7% worked fully in person. In terms of their immediate work environment, nearly half of all participants reported being the only scientific editor in their workgroup (46%), even among those who were institutionally employed. Being a freelancer or remote worker was associated with being the only scientific editor in a group, with 96% of freelancers and 55% of institutionally employed but fully remote employees reporting working alone.

Participants’ most common annual salary was $75,000-$100,000 (29%), followed closely by $100,000-$125,000 (24%) and $50,000-$75,000 (21%; Table 3). Only 8% of participants reported earning $0-$30,000 per year. Of those who reported an annual salary of less than $50,000, all identified as being freelancers or part-time employees. Possession of a PhD was associated with having a higher salary. PhD recipients had a mean salary of $105,400, while those with a master’s degree had a mean salary of $79,700, and BA/BS $78,400.

**Table 3.** Reported Annual Salaries for Scientific Editors in 2024. Bar chart is normalized to the largest value.

| Salary range | Percent of participants |
| --- | --- |
| \$0-\$30,000* | 8% |
| \$30,000-\$50,000* | 3% |
| \$50,000-\$75,000 | 21% |
| \$75,000-\$100,000 | 29% |
| \$100,000-\$125,000 | 24% |
| \$125,000-\$150,000 | 11% |
| \$150,000-\$175,000 | 4% |
| \$175,000-\$200,000 | 4% |
| \$200,000+ | 1% |

\*All survey participants in the \$0-\$30,000 and \$30,000-\$50,000 categories also described their employment as part-time or freelance.

The main client base for scientific editors included *Faculty* (80%), *Research staff* (61%), *Trainees* (60%), and/or *Administrators* (31%). *Other* responses (19%) included medical writers, large institutions or agencies, journals and publishers, and other health care professionals (e.g., nurses). Twenty percent of survey participants edited for all four groups (*Faculty*, *Research staff*, *Trainees*, and *Other*). Many participants (61%) stated that they edit for 30 or fewer clients per year, with the most common response (22%) being 11-20. Four percent of participants edited for more than 100 clients.

### Career paths and defining success

To identify career paths that lead to becoming a scientific editor, we asked participants to describe their prior work experience. Responses included a total of 28 different job classifications, most of which could be categorized into one of three larger groups: higher education, medical/clinical settings, or publishing. Specific positions previously held by survey participants included scientist (53%), copy editor (44%), instructor or faculty member (38%, combined), freelance writer (26%), and to a lesser extent, more varied alternatives, such as journal editors (15%), grant coordinators (9%), and business managers (5%).

Participants entered the scientific editing field at various stages of their education or career paths, ranging from *after undergraduate studies* to *after being a faculty member*, with a large number (65%) beginning sometime after graduate studies (Table 4). However, a small number of participants (6%) began editing after working in an unrelated career.

**Table 4.**
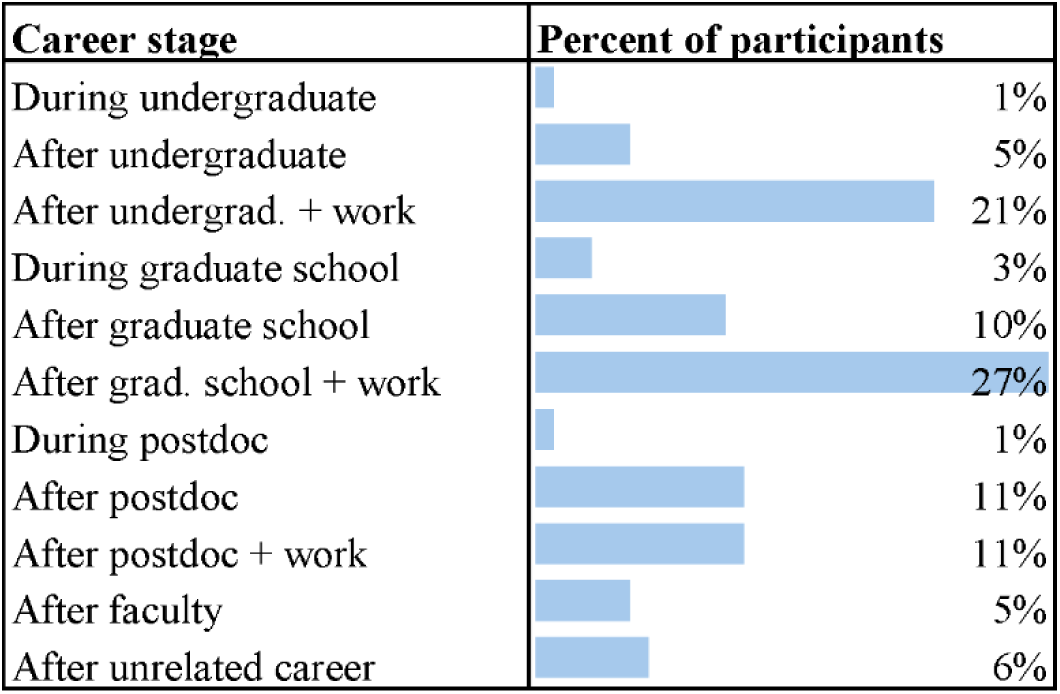
Career Stage when Participants Entered the Field of Scientific Editing. Bar chart is normalized to the largest value.

The most common response to the question of how scientific editors learned of the profession was by doing informal work for colleagues (27%). Another 24% of participants learned about the profession from a friend, colleague, or mentor. Only 22% of participants intentionally sought a position as a scientific editor or related occupation. *Other* responses (27%) included being introduced to the profession in a variety of ways: job ads or fairs, reading or seeing presentations about scientific editing, or gradually moving into an editing position as part of other responsibilities. Only one participant was advised by a colleague or career counselor to go into scientific editing, and one other learned about the profession through an internship related to scientific editing.

Despite the inconsistencies in career pathways and professional training, participants most commonly acquired their first scientific editing position (54%) through posted job opportunities. A number of people simply began independent freelancing (18%), and others were hired into the position through a friend or colleague (15%). Some began informal editing activities for colleagues, which expanded into a formal position (9%). *Other* means of acquiring the position, such as recruitment, evolution of job duties, internships, and a cold approach to an employer, were reported by 7% or less of participants.

The survey queried how long participants had been in the scientific editing profession. The smallest percentage had been hired in the previous five years (12%, combined categories), with increasing proportions of experience among the remaining participants. Well over half (68%) had been hired more than eleven years previously, and 28% had been in the profession more than 20 years.

We asked participants about available career advancement pathways with their current employer. The most common pathways included a manager/director track (34%) and a scientific editor track (31%). Other possibilities for career development included an administrator track (13%); an editorial track, such as at a journal (6%); and a faculty track (6%). Twenty-six percent of participants reported they work as freelancers (this number includes those who freelanced exclusively as well as those who freelanced in addition to a regular position), 14% reported that they are unsure of career advancement options, and 13% reported not having a pathway for career advancement. Notably, among institutionally employed participants, those who worked in a larger office were significantly more likely to report having a clear path for advancement than those who worked alone (80% versus 55%).

To determine how participants assess the impact of their work, we asked if and how they track outcomes. More than half (64%) reported that they do. Some of the methods reported for tracking outcomes included client feedback through surveys or inquiries, results from online database searches of publications and/or funded grants; and outcomes as tracked by a department. Client satisfaction (71%) was the most common measure participants sought to assess. Other measures of impact included the success rate of submitted grants or manuscripts (58%), feedback received by reviewers (8%), or the number of submitted manuscripts and proposals worked on (6%). Other means for measuring success included service volume and activity, internal evaluations and standards, value added, and trainings developed and delivered.

### Time spent editing

We asked participants to list the percentage of time spent on various activities, with time totaling 100%. All participants reported spending time *Editing* (Table 5A), with the two most common ranges being 41-50% of time (21% of participants) and 71-80% of time (20% of respondents) (Table 5B). Only 1% of participants spent less than 10% of their time editing, whereas 10% reported spending 91-100% of their time editing (Table 5B).

**Table 5.** Activities Performed by Scientific Editors and Amount of Time Spent on Editing. (A) Percentage of participants who reported partaking in the indicated activities. (B) Percentage of time editors reported spending on editing. Bar charts in (A) and (B) are normalized to the largest value in the respective category.

| A) % of participants who perform selected activities |  | B) % of time spent on editing |  |
| --- | --- | --- | --- |
| Editing | 100% | ≤10% | 1% |
| Administrative work | 73% | 11-20% | 4% |
| Mentoring | 46% | 21-30% | 8% |
| Teaching | 38% | 31-40% | 11% |
| Communications | 36% | 41-50% | 21% |
| Other | 35% | 51-60% | 10% |
| Management/supervision | 25% | 61-70% | 8% |
| Grant coordination | 21% | 71-80% | 20% |
| Research assistance | 15% | 81-90% | 8% |
| Publication management | 11% | 91-100% | 10% |

In addition to editing, participants reported spending time on administrative work, mentoring, teaching, communications, management/supervision, grant coordination, research assistance, and publication management (Table 5A). Among participants who selected *Other*, activities included grant writing, funding searches, research into journal guidelines, grant library management, professional development, visuals (graphic design), grant planning meetings, writing, presentations, and helping investigators find resources.

### Types of editing provided

We surveyed participants on the types of editing they do, as well as content areas and types of documents on which they worked. Nearly all participants performed substantive editing (97%), and almost as many also edited for grammar/mechanics (93%). Over half of participants reported assisting with strategy/grantsmanship (58%). Forty-five percent reported editing for compliance, and 45% indicated they provide feedback on the science content. *Other* responses included editing to improve the style/format of documents and fact checking.

In terms of subject matter, most participants stated that they edit documents in the biomedical sciences (85%), biology (57%), and/or public health (57%). Approximately a quarter of participants indicated they edit in the areas of engineering (26%), chemistry (24%), or the humanities (24%); 18% edit physics documents and 28% of participants indicated they edit all the content areas that were listed.

In terms of document types, participants reported that they edit manuscripts and scientific publications (84%), grant/fellowship proposals (75%), and materials for lay audiences (52%); 3% edited all three types of documents. Participants also reported editing white papers (41%) and FDA or regulatory documents (9%). *Other* document types were edited by 12% of participants, with examples including abstracts, posters, and presentations; academic content (e.g., dissertations, educational content), career-related documents (e.g., CVs, letters of recommendation), procedures, and books.

### Skills needed for scientific editing

We asked participants to list the skills they felt were important to be successful at scientific editing. The most commonly reported skills were efficient work methods (55%), interpersonal/client relations skills (49%), language/writing skills (46%), emotional intelligence (35%), subject matter expertise (35%), and critical thinking skills (25%). Specific characteristics within each category are listed in Table 6.

**Table 6.**
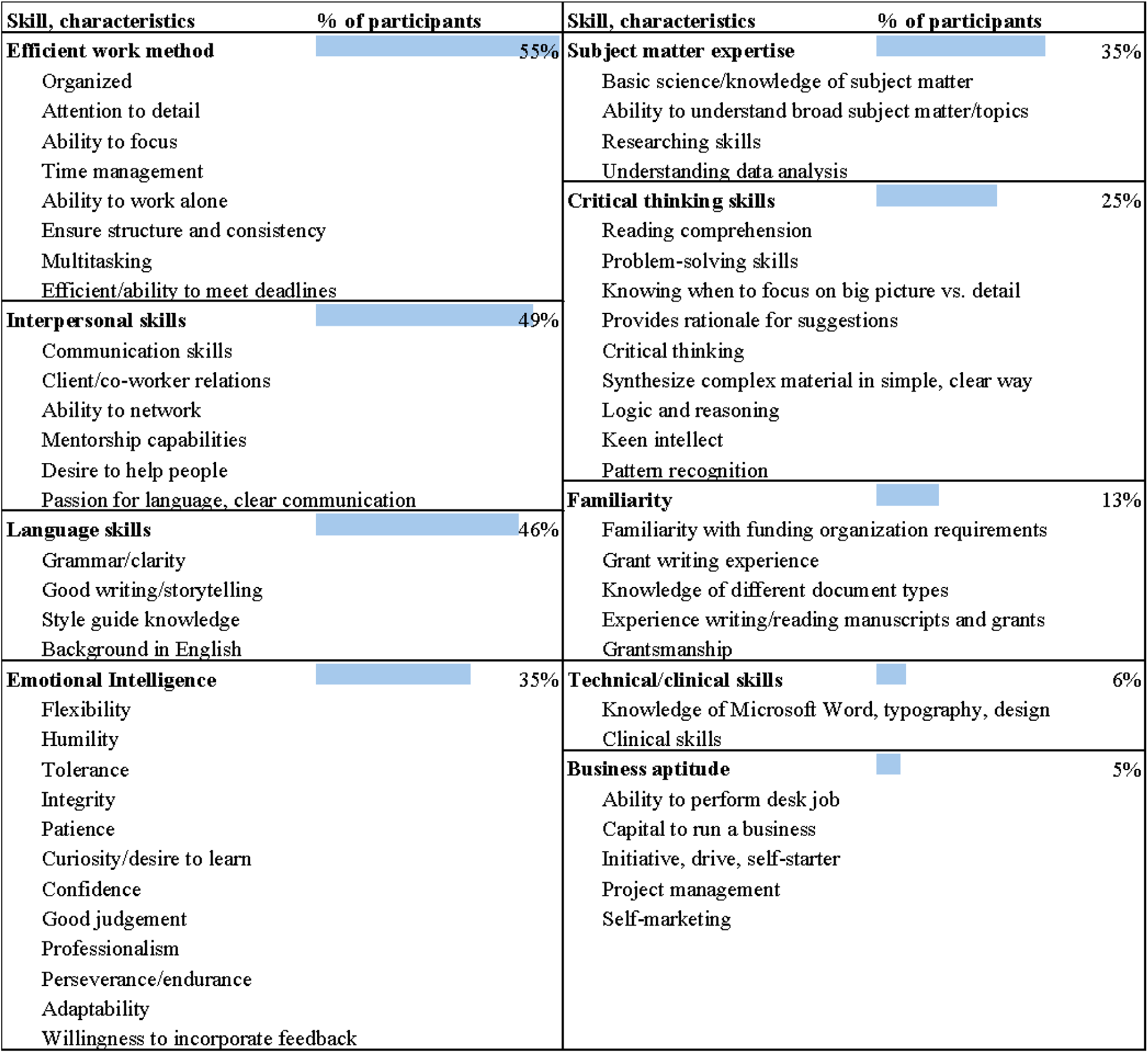
Important Skills of Scientific Editors. Participants were asked to list the three skills they feel are most important for work as a scientific editor. Skills were grouped into nine categories, and individual characteristics within each category are listed. Bar charts are normalized to the largest value of all categories.

| Skill, characteristics | % of participants | Skill, characteristics | % of participants |
| --- | --- | --- | --- |
| **Efficient work method** | 55% | **Subject matter expertise** | 35% |
| Organized |  | Basic science/knowledge of subject matter |  |
| Attention to detail |  | Ability to understand broad subject matter/topics |  |
| Ability to focus |  | Researching skills |  |
| Time management |  | Understanding data analysis |  |
| Ability to work alone |  | **Critical thinking skills** | 25% |
| Ensure structure and consistency |  | Reading comprehension |  |
| Multitasking |  | Problem-solving skills |  |
| Efficient/ability to meet deadlines |  | Knowing when to focus on big picture vs. detail |  |
| **Interpersonal skills** | 49% | Provides rationale for suggestions |  |
| Communication skills |  | Critical thinking |  |
| Client/co-worker relations |  | Synthesize complex material in simple, clear way |  |
| Ability to network |  | Logic and reasoning |  |
| Mentorship capabilities |  | Keen intellect |  |
| Desire to help people |  | Pattern recognition |  |
| Passion for language, clear communication |  | **Familiarity** | 13% |
| **Language skills** | 46% | Familiarity with funding organization requirements |  |
| Grammar/clarity |  | Grant writing experience |  |
| Good writing/storytelling |  | Knowledge of different document types |  |
| Style guide knowledge |  | Experience writing/reading manuscripts and grants |  |
| Background in English |  | Grantsmanship |  |
| **Emotional Intelligence** | 35% | **Technical/clinical skills** | 6% |
| Flexibility |  | Knowledge of Microsoft Word, typography, design |  |
| Humility |  | Clinical skills |  |
| Tolerance |  | **Business aptitude** | 5% |
| Integrity |  | Ability to perform desk job |  |
| Patience |  | Capital to run a business |  |
| Curiosity/desire to learn |  | Initiative, drive, self-starter |  |
| Confidence |  | Project management |  |
| Good judgement |  | Self-marketing |  |
| Professionalism |  |  |  |
| Perseverance/endurance |  |  |  |
| Adaptability |  |  |  |
| Willingness to incorporate feedback |  |  |  |

### Professional resources and support

We asked what resources editors use daily and found 67% of participants rely heavily on journal or sponsor websites, 61% on style guides, 38% on templates, and 22% on books or workbooks. Another 19% listed PubMed, Google Scholar, Wikipedia, and in-house materials among *Other* resources. Most editors reported using several different types of resources on a daily basis.

Scientific editors employed by a university, research institution, or other organization (*n*=88) reported a variety of employer-provided resources to facilitate their work. These resources included subscriptions to reference books, journals, or other online resources (74%); access to written guidelines or style guides (70%); access to different software packages (70%); membership in professional societies (63%); internal presentations or workshops (61%); funding to attend local or national conferences (61%); and *Other* resources (10%), which included specific trainings or online resources and professional communities within their institutions. Participants who self-identified exclusively as self-employed or freelancers did not have employer-provided resources, and were thus excluded from this survey question.

Survey participants were asked to name any employer-provided resources that would have been useful when they began work as scientific editors, as well as any that they wished were available now (Table 7). Among these participants, 23% reported that training from workshops and webinars would have benefited them early in their careers, while none reported that they currently wished for access to workshops and webinars, indicating fulfillment of that need. Similarly, 19% reported that direct training through personal mentorship or exposure to the profession was lacking early in their career, while only 2% reported the same need currently. Twelve percent reported a need for formalized coursework or certifications related to the profession early in the career, compared with just 5% who reported a current need for this resource. Conversely, no participants reported that they wished for access to journals and online/software tools early in their editorial careers, while 15% reported a desire for greater access to these resources currently, perhaps indicating a later realization of that need. Participants reported similar needs for guidebooks, style guides, and other guideline literature (11% early in the career, 16% currently); networking and conferences (17% early in the career, 15% currently); and membership in a professional organization (7% early in the career, 4% currently) between their early and current careers. The percentage of participants satisfied with available resources was 22% early in their career and 34% currently.

**Table 7.** Professional Resources in Demand. Participants reported which resources they wish had been available to them early in their careers as scientific editors (A), and which professional resources they wished to have currently (B). Bar charts in (A) and (B) are normalized to the largest value in each respective category.

| Resource | A) Early career, % | B) Currently, % |
| --- | --- | --- |
| Workshops and webinars | 23% | 0% |
| Exposure, mentorship, training | 19% | 2% |
| Networking and conferences | 17% | 15% |
| Coursework or certifications | 12% | 5% |
| Literature and style guides | 11% | 16% |
| Professional organizations | 7% | 4% |
| Journal access, software, and online tools | 0% | 15% |
| Other | 8% | 15% |
| Satisfied/NA | 22% | 34% |
| Unsure | 2% | 4% |

When asked to name their favorite source for professional development or support, most participants cited one or more professional organizations (76%), including the National Organization of Research Development Professionals (NORDP; 20%), Scientific Editors Network (ScENe; 20%), American Medical Writers Association (AMWA; 18%), and Board of Editors in the Life Sciences (BELS; 14%). *Other* responses were categorized as various online resources (9%), the support of colleagues (7%), and internal office or organizational resources (4%). It should be noted that many of the professional organizations were also the primary means for disseminating the survey, and many editors belonged to more than one organization.

### Challenges and rewards of scientific editing

We asked participants to list their three least favorite and three most favorite aspects of being a scientific editor. Almost half of participants (48%) indicated dealing with difficult clients (e.g., those who were overly demanding, did not communicate well, behaved disrespectfully, or requested services too late to enable quality editing) as their least favorite aspect of being a scientific editor. Issues related to timing (38%; e.g., working nights or weekends, being on-call around grant deadlines, rushing through tight timelines) represented the second most common disliked aspect, followed by challenges with the type of editing project (36%; multiple authors, unfamiliar subject) and feeling undervalued/lack of advancement (24%).

In response to the question about their three favorite aspects of being a scientific editor, two-thirds of participants (67%) cited the intellectually interesting nature of the profession. Other favorite aspects included the satisfaction of helping others (48%), the opportunity to work with educated people (i.e., faculty, researchers, editors; 42%), work-life flexibility (35%), and making a meaningful contribution to science (29%).

## Discussion and Conclusions

One of the primary themes from the survey was that scientific editors have homogeneous demographic characteristics. Our findings (83% female and 87% White) are consistent with the demographics of research development professionals overall (82% female and 83% White, per the 2025 NORDP survey).^17^ Respondents were also highly educated (43% PhD and 26% master’s degree), with most having an educational background in biomedical or natural sciences (39% and 27%, respectively). The majority of undergraduate and postgraduate degree recipients, including recipients of PhDs in life sciences, are White and female; thus, our findings seem to also reflect broader U.S. educational trends.^23^ Black and Latino employees are significantly underrepresented in STEM fields overall, which aligns with our findings (1% and 3% of respondents, respectively).^24^ One explanation for the overrepresentation of women in our survey is that despite their educational attainment, women leave academic medicine at higher rates than do men,^25^ and scientific editing (and research development in general) is a natural career transition. Indeed, 53% of our respondents indicated that they had previous work experience as a scientist, with 38% having served as an instructor or faculty member. Our findings provide a useful foundation for research into the underlying causes of disparities in gender and race/ethnicity in scientific editing, which we aim to further explore in our follow-up survey.

### Career Path

Our data show that scientific editors come from a variety of backgrounds and experiences. The wide variability in the career stage at which participants began editing professionally (Table 4) suggests that most people enter into scientific editing positions as a result of filling a need or expanding other responsibilities or interests as opposed to a deliberate pursuit of a position at the early career stage. Even the Occupational Outlook Handbook, used by high school career advisors since the post-WWII era, does not have entries for *science writer*, *science editor*, or even *grant writer*. The closest job category is that of *technical writer*,^19^ the duties of which are described as “[to] prepare instruction manuals, how-to guides, journal articles, and other supporting documents to communicate complex and technical information more easily.” According to the handbook, the job “typically” requires a bachelor’s degree, while acknowledging that “knowledge of or experience with a technical subject, such as science or engineering, is beneficial.”^26^ Although scientific editing is rarely named as a distinct formal career path, many of the competencies of scientific editors overlap with those of research development professionals, especially assisting with grant proposal development, offering grantsmanship support, and providing strategies to enhance the competitiveness of research proposals or manuscripts. Our survey responses clearly demonstrate a lack of composite descriptions of job preparation, classification, or title: 15 participants held job titles that explicitly mentioned “research development”, and an additional 31 held titles that did not mention research development but were NORDP members who performed essentially the same duties as research development professionals. This combined group of 46 participants was more likely to hold a PhD. They also predominantly edited grants versus other types of documents.

Holland’s typology of career choices offers a theory that may contribute to understanding the disparate backgrounds of scientific editors as well as the larger field of research development professionals.^27^ Simply put, Holland argued that people flourish in working environments that are congruent with their personality type. While our survey did not overtly attempt to uncover respondents’ personality features, the question regarding favorite job aspects offers some clues: scientific editors generally appreciate intellectually interesting work, helping others, and work-life flexibility. Scientific editing offers all of these elements, perhaps in a unique formulation.

Since scientific editors also skew toward high educational levels, intellectual challenge is a logical occupational congruency. Yet the question of editors’ disparate backgrounds might best be pursued by investigating the push factors from their previous occupations rather than the pull factors that drew them into scientific editing. According to Holland’s theory, the favorite job aspects reported in our survey may be a reflection that these characteristics were lacking for scientific editors in their previous positions yet rather rich in scientific editing. As with other survey results, the diversity of career paths into scientific editing has prompted more questions than answers. A future survey could pursue questions of job satisfaction and persistence in the profession.

### Training and Professional Development

Training and professional preparation varied widely in our survey, with no common curriculum or training program for becoming a scientific editor being apparent. Although universities such as Johns Hopkins and Temple University offer master’s degrees in science writing, no participants indicated such a degree. There are also scattered undergraduate minors and certificates in U.S. universities, as well as certifications offered by professional organizations such as BELS. Research development is included in some broader career development programs offered by many universities, and scientific editing is included in the limited number of research development internship programs available. Scientific editing professionals are often expected to have at least an undergraduate major in a scientific field. Our data show that while many participants hold advanced and professional degrees, they collectively entered the scientific editing profession at almost every possible phase of their education or career, ranging from those who started editing as an undergraduate student to those who began editing while in a faculty position. The inconsistency in educational attainment may help explain the diversity of pathways to the career, and perhaps why career advancement as a scientific editor can be nebulous.

The participants in this survey largely represented either individuals employed by higher education and medical research institutions or freelancers whose clients generally work for similar science institutions. Participants worked in almost all academic disciplines, from biomedical sciences to humanities to computer science. They conducted a myriad of activities and were generally satisfied with their work and career. Career track pathways also varied, and over a quarter of participants had no formal career advancement pathway or were unsure about available career advancement options.

Scientific editing has long been connected to well-established professional organizations, such as AMWA (founded in 1940)^28^ and BELS (founded in 1991).^29^ In academic settings, scientific editors are often closely associated with the research development profession (NORDP; founded in 2010).^30^ Our survey results indicated widespread commitment to professional memberships among participants. The question regarding such affiliations yielded a total of 300 organizational affiliations from 127 participants, an average of 2.36 memberships per person. Only three participants indicated that they did not hold membership in any professional group. Interestingly, NORDP members were likely to be ScENe members. We also observed that joint membership in BELS and AMWA was common. While many people reported these organizations as important sources of training and professional development, the rate of membership among scientific editors suggests these organizations also offer a sense of community that members appreciate and support.

While many survey participants did not necessarily set out to be scientific editors, they were collectively quite experienced and proficient in their jobs. Well over half of participants (68%) were hired more than eleven years prior to the survey, and over one-quarter (28%) reported having been in the profession for more than 20 years. The survey results indirectly showed that employment trends and entry into the profession seem to coincide with national economic trends, apparently as higher education institutions tighten or loosen their overall budgets. For example, fewer scientific editors started their jobs 16-20 years previously (15%) as compared to other five-year increments. The time frame of 2004-2009 coincides with the great recession, an era during which numerous states reduced their appropriations on colleges and universities.^31,32^ Higher education has been undergoing transition since the 1980s in a myriad of ways, not the least of which has been a declining share of resources going to instruction.

Administrative hiring across the country is a significant part of that transfer of resources. However, beginning in 2008 during the recession, numerous institutions sought to reduce administrative costs as part of their restructuring plans.^33^ Scientific editing positions may have been part of those reductions. Non-academic staff hiring increased after 2012, in areas focused on regulatory compliance, information technology, and fundraising and development.^34^ Scientific editing, like research development, is tasked with helping institutions win grants and thus can be considered an investment in institutional development.

Twelve percent of survey participants entered the profession in the previous five years. While this statistic might reflect a bias in survey response (i.e., those newer to the profession did not respond as readily as others or do not yet belong to any of the targeted organizations), it is also possible that the results reflect fewer people entering the scientific editing profession during the last few years.

As one might expect, survey data indicate that a background in science is valued in this profession, perhaps more than a background in English or journalism, although there is a merging of skillsets from both sectors. Furthermore, the data in Table 6 indicate that scientific editing requires several other skillsets beyond science and language, including interpersonal, organizational, and critical thinking skills. However, our data also demonstrated a lack of a clear career path for scientific editors. This may represent an opportunity for the growth or expansion of research development offices at universities and research organizations, which are well-positioned to incorporate scientific editors given the overlap in the functions performed by each.

### Limitations

Since scientific editors often belong to two or more professional organizations, and the individual share rate is unknown, we do not know how many unique individuals received the invitation to the survey. Additionally, because we recruited survey participants from national organizations, our results are mostly limited to memberships in those organizations and may not reflect all scientific editors in the profession. This survey did not include journalistic science writers who work for news agencies or popular/social media, although some of their skills and experience may overlap. An additional limitation relates to the potential overlap in professional roles captured in our study. While we took steps to evaluate survey validity during the development process, some participants could have interpreted questions differently based on their role and experience. Specifically, because of the interdisciplinary and evolving nature of scientific editors and research development professionals, and the reliance on individuals to categorize their own work as “scientific editing,” there may some ambiguity in the interpretation of our findings. Thus, a follow-up survey could use more discrete criteria for scientific editors to enable distinction between these two roles. Moreover, research development is a growing field in the U.S., but it is not as well-defined in other countries. Therefore, the data are specific to those working in the U.S. and the implications of our findings are not generalizable for global research development practice.

The ScENe survey was completed in 2024, before the widespread adoption of large language model artificial intelligence tools. Therefore, our survey did not include any questions regarding these tools. In our follow-up survey, we will explore how the increasing use of generative and non-generative artificial intelligence tools have affected the scientific editing field (e.g., whether and how respondents use such tools, how these tools have affected the quality or quantity of documents they edit, and whether they believe the increased use of these tools has affected the demand for professional editors). The survey also took place after the COVID-19 pandemic subsided but before significant changes in the federal science funding landscape took place in 2025. While the survey seems to reflect a valid characterization of the field in this time frame, which is a valuable data point, the larger world of academic and grant proposal writing and editing is likely to shift significantly in the future. Thus, we expect a follow-up survey, in or around 2030, will likely yield substantially different data, perhaps demonstrating a changed workforce in terms of numbers, responsibilities, methods, and standards. The rapidly changing technology and the research environment may preclude some companies and institutions from hiring or consulting scientific writers and editors. With many grants revoked, programs cancelled, and overhead rates potentially reduced, many higher education institutions will presumably be in retraction mode for the foreseeable future. On the other hand, research grants and publishing will become increasingly competitive, and scientific editors proportionately more valuable to their clients.

## Conclusions

By synthesizing the findings of the 2024 ScENe Survey and related literature, this article provides a comprehensive overview of the state of the scientific editing profession, highlights its importance to the research enterprise, and characterizes its integration within the research development profession. It also provides a dataset that can be further interrogated for associations or comparisons with other related studies. Given the overlap in the activities performed by scientific editors and research development professionals, and the lack of clearly defined career pathways for scientific editors within research development offices, our findings suggest an opportunity to expand the scope of these offices, either by integrating scientific editors into existing workflows or by establishing dedicated career pathways for them.

Accordingly, this study highlights directions for future research, professional development, and institutional assessment to support the integration of scientific editors within research development offices. Because our survey was conducted prior to the rapid emergence of artificial intelligence tools, including large language models, it also provides a valuable benchmark to track progress and change within the field as more data are gathered in future surveys.

## Acknowledgements

The ScENe manuscript committee would like to thank ScENe leadership for their support and advice. We also thank our survey beta testers for their helpful feedback.

## List of Supplementary Materials

Table S1. Full Survey Data. Survey question text, values for responses, distinction for responses selected from multiple choices or those categorized by inductive coding, and number of respondents to individual questions (n).

