## Supplemental Table 1 for "Careers in Scientific Editing: Results from the 2024 Scientific Editors Network (ScENe) Survey"

***Table S1.*** *Full Survey Data. Survey question text, values for responses, distinction for responses selected from multiple choices or those categorized by inductive coding, and number of respondents to individual questions (n).*

| **Q.1: Which of the following activities do you perform as part of your job? Please select all that apply.** Selected choices. (*n*=145) | |
|---|---|
| Scientific editing of research grants, manuscripts, or other scholarly works | 90% (131) |
| Researching grant-writing strategies (e.g., grantsmanship) for scientific/research purposes | 39% (56) |
| Providing instruction in scientific writing (e.g., teaching, seminars, mentorship) | 30% (44) |
| None of these | 6% (9) |

| **Q.2: Please select your age group (years).** Selected choices. (*n*=132) | |
|---|---|
| 18-24 | 0% (0) |
| 25-34 | 9% (12) |
| 35-44 | 23% (31) |
| 45-54 | 30% (40) |
| 55-64 | 27% (36) |
| 65+ | 10% (13) |
| Prefer not to answer | 0% (0) |

| **Q.3: What gender do you identify as?** Selected choices. (*n*=132) | |
|---|---|
| Female | 83% (109) |
| Male | 17% (22) |
| Non-binary | 0% (0) |
| Other | 0% (0) |
| Prefer not to answer | 1% (1) |

| **Q.4: What is your primary country of residence (and region, if US)?** Selected choices. (*n*=132) | |
|---|---|
| United States of America: Northeast (Connecticut, Maine, Massachusetts, New Hampshire, Rhode Island, Vermont) | 9% (12) |
| United States of America: Atlantic (Delaware, Maryland, New Jersey, New York, Pennsylvania, District of Columbia, Virginia) | 18% (24) |
| United States of America: Southeast (Alabama, Arkansas, Florida, Georgia, Kentucky, Louisiana, Mississippi, North Carolina, Puerto Rico, South Carolina, Tennessee, U.S. Virgin Islands, West Virginia) | 17% (22) |
| United States of America: Great Lakes (Illinois, Indiana, Michigan, Minnesota, Ohio, Wisconsin) | 11% (15) |
| United States of America: Midwest/Mountain (Iowa, Kansas, Missouri, Nebraska, North Dakota, South Dakota, Idaho, Montana, Wyoming, Nevada, Colorado, Utah) | 12% (16) |

| | |
|---|---|
| United States of America: Southwest (Arizona, New Mexico, Oklahoma, Texas) | 14% (18) |
| United States of America: Pacific (Alaska, California, Hawaii, Oregon, Washington) | 11% (14) |
| Canada | 3% (4) |
| Other | 5% (7) |
| Prefer not to answer | 0% (0) |

1

| **Q.5: What is your race or ethnicity? Please select all that apply.** Selected choices. (*n*=132) | |
|---|---|
| American Indian or Alaskan Native | 0% (0) |
| Asian | 11% (15) |
| Black or African American | 1% (1) |
| Hispanic or Latino | 3% (4) |
| Native Hawaiian or Pacific Islander | 0% (0) |
| White | 87% (115) |
| Other | 2% (2) |
| Prefer not to answer | 1% (1) |

2

| **Q.6: Is English your native language?** Selected choices. (*n*=132) | |
|---|---|
| Yes | 95% (125) |
| No | 5% (7) |

3

| **Q.7: What is the highest level of education you have completed? Please select all that apply.** Selected choices. (*n*=132) | |
|---|---|
| High school | 3% (4) |
| Associates degree | 2% (2) |
| BA/BS | 26% (34) |
| MA/MS | 26% (34) |
| MPH | 2% (3) |
| PhD | 43% (57) |
| MD/DO | 0% (0) |
| MBA | 1% (1) |
| Other | 6% (8) |

4

| **Q.8: What is your education background? Please select all that apply.** Selected choices. (*n*=132) | |
|---|---|
| Biomedical Sciences | 39% (52) |
| Natural Sciences | 27% (36) |
| English | 21% (28) |
| Communications or Journalism | 17% (22) |
| Public Health | 5% (6) |
| Medicine | 2% (3) |
| Library and Information Science | 2% (2) |

| | |
|---|---|
| Other | 27% (35) |

| **Q.9a: In addition to your educational background, what kind of formal training have you had in scientific editing? Please select all that apply.** Selected choices. (*n*=131) | |
|---|---|
| Scientific editing internship (see Q.9b-9g) | 4% (5) |
| Scientific editing courses/workshops | 44% (58) |
| Mentoring in scientific editing | 28% (37) |
| Grant administration | 14% (19) |
| Other | 23% (31) |
| None | 26% (34) |

| **Q.9b: How did you hear about the internship?** Categorized responses. (*n*=4) | |
|---|---|
| Training position | 25% (1) |
| Advertised to students (campus newspaper, department, instructor, etc.) | 75% (3) |

| **Q.9c: What type of employer provided the internship?** Selected choices. (*n*=4) | |
|---|---|
| University | 25% (1) |
| Private company | 0% (0) |
| Professional organization | 0% (0) |
| Publisher | 50% (2) |
| Other | 25% (1) |

| **Q.9d: How long was the internship?** Selected choices. (*n*=4) | |
|---|---|
| <6 months | 0% (0) |
| 6 months-1 year | 50% (2) |
| 1-2 years | 50% (2) |
| 2+ years | 0% (0) |
| Other | 0% (0) |

| **Q.9e: What tasks did you perform as part of your internship? Please select all that apply**. Selected choices. (*n*=4) | |
|---|---|
| Editing | 100% (4) |
| Reviewing relevant materials/resources | 75% (3) |
| Teaching | 25% (1) |
| Consulting | 25% (1) |
| Help staff develop content | 50% (2) |
| Other | 0% (0) |

| **Q.9f: What knowledge and skills did you gain in your internship?** Categorized responses. (*n*=4) | |
|---|---|
| Scientific/technical editing experience | 100% (4) |
| Leading discussions with corresponding investigators or clinicians | 50% (2) |
| Journal submission processes | 25% (1) |

| | |
|---|---|
| Grantsmanship | 25% (1) |
| Providing feedback (written and/or verbal) | 25% (1) |

1

| **Q.9g: On a scale from 1-5 how well do you feel your internship prepared you for work as a scientific editor?** (*n*=4) | |
|---|---|
| 1 | 25% (1) |
| 2 | 0% (0) |
| 3 | 0% (0) |
| 4 | 25% (1) |
| 5 | 50% (2) |

2

| **Q.10: What prior work experience do you bring to your work as a scientific editor? Please select all that apply.** Selected choices. (*n*=129) | |
|---|---|
| Scientist | 53% (68) |
| Copy Editor | 44% (57) |
| Freelance Writer | 26% (33) |
| Instructor | 24% (31) |
| Research Development Officer | 16% (21) |
| Communications/Marketing Specialist | 16% (20) |
| Journal Editor | 15% (19) |
| Research Administrator | 15% (19) |
| Faculty member | 14% (18) |
| Journalist | 10% (13) |
| Grant Coordinator | 9% (11) |
| Administrator at a journal | 6% (8) |
| Other writing/editing | 6% (8) |
| Business Manager/Director/Coordinator | 5% (7) |
| Clinician or Allied Health Professional | 5% (7) |
| Other Education or Technology-Related | 5% (6) |
| None of the above | 2% (2) |

3

| **Q.11: What professional certifications related to scientific writing/editing do you have? Please select all that apply.** Selected choices. (*n*=131) | | |
|---|---|---|
| Editor in the Life Sciences (ELS) | | 46% (60) |
| AMWA Certificate (Essential Skills) | | 11% (14) |
| Diplomate Editor in the Life Sciences (ELS[D]) | | 5% (6) |
| Medical Writer Certified (MWC) | | 2% (2) |
| EMWA Certificate [Basic and Advanced] | | 1% (1) |
| None | | 47% (61) |
| Other (please specify) | | 8% (10) |
| AMWA Certificate [Basic and Advanced] | 4% (5) | |
| Certified Research Administrator (CRA) | 1% (1) | |

| | |
|---|---|
| Society for Technical Communication, Certified Professional Technical Communicator | 1% (1) |
| Institution-specific | 1% (1) |
| Non-U.S. (English) | 2% (2) |

| **Q.12: What is your current employment status?** Selected choices. (*n*=127) | |
|---|---|
| Employed full-time | 70% (89) |
| Employed part-time | 2% (2) |
| Freelance | 20% (26) |
| Retired | 2% (3) |
| Other | 6% (7) |

| **Q.13: What type of institution do you work at?** Selected choices. (*n*=84) | |
|---|---|
| Public | 57% (48) |
| Private | 43% (36) |

| **Q.14: Where are you currently employed? Please select all that apply.** Selected choices. (*n*=121) | |
|---|---|
| University | 40% (49) |
| I am a freelancer | 29% (35) |
| Research institution | 18% (22) |
| Hospital | 16% (19) |
| Medical college | 12% (14) |
| Non-profit organization | 8% (10) |
| Medical communications/marketing company | 7% (8) |
| Publisher | 2% (3) |
| Pharmaceutical company | 2% (2) |
| Professional society/organization | 2% (2) |
| Other | 8% (10) |

| **Q.15: What is your work arrangement?** Selected choices. (*n*=122) | |
|---|---|
| Fully remote (includes freelancers) | 64% (78) |
| Hybrid | 30% (36) |
| Fully in person | 7% (8) |

| **Q.16: Who do you edit for? Please select all that apply.** Selected choices. (*n*=119) | | |
|---|---|---|
| Trainees (undergraduate and graduate students, postdocs, fellows, residents) | | 60% (71) |
| Research staff | | 61% (72) |
| Faculty | | 80% (95) |
| Administrators | | 31% (37) |
| Other (please specify) | | 19% (23) |
| Uncategorized | 6% (7) | |

| | |
|---|---|
| Medical writers | 5% (6) |
| Large institution (e.g., professional associations, pharmaceutical companies) | 3% (4) |
| Publisher/journal | 3% (3) |
| Healthcare provider (e.g., nursing, private practice) | 3% (3) |

1

| **Q.17: What is your current job title?** Categorized responses. (*n*=122) | |
|---|---|
| Title includes: “Editor” | 49% (60) |
| Title includes: “Writer” | 16% (20) |
| Title includes: “Research Development” | 8% (10) |
| Title includes: “Grant” | 7% (9) |
| Title includes: “Research” (but not “Development”) | 12% (15) |
| Title includes: “Science” or “Scientific” | 27% (33) |
| Title includes designation of level (junior, associate, etc.) | 35% (43) |
| Title includes multiple terms | 12% (14) |

2

| **Q.18: What is your current annual salary?** Selected choices. (*n*=111) | |
|---|---|
| $0-$30,000 | 8% (9) |
| $30,000-$50,000 | 3% (3) |
| $50,000-$75,000 | 21% (23) |
| $75,000-$100,000 | 29% (32) |
| $100,000-$125,000 | 24% (27) |
| $125,000-$150,000 | 11% (12) |
| $150,000-$175,000 | 4% (4) |
| $175,000-$200,000 | 4% (4) |
| $200,000+ | 1% (1) |

3

| **Q.19: Including yourself, how many full-time equivalent scientific editors work in your office/department?** Categorized responses. (*n*=120) | |
|---|---|
| Nobody is full-time in the office | 6% (7) |
| I’m all alone | 46% (55) |
| Me and someone else | 13% (16) |
| 3-5 people | 14% (17) |
| 6-10 people | 8% (9) |
| 11-20 people | 8% (9) |
| 20+ people | 3% (3) |
| Unknown | 3% (4) |

4

| **Q.20: What types of documents do you edit? Please select all that apply.** Selected choices. (*n*=119) | | |
|---|---|---|
| Manuscripts and scientific publications | | 84% (100) |
| Grant/fellowship proposals | | 75% (89) |
| Materials for lay audience | | 52% (62) |

| | | |
|---|---|---|
| White papers | | 41% (49) |
| FDA submissions or regulatory documents | | 9% (11) |
| Other (please specify) | | 12% (14) |
| Abstracts, posters, presentations | 4% (5) | |
| Academic (e.g., theses) | 3% (4) | |
| Career (e.g., CVs, cover letters) | 3% (4) | |
| Procedures | 3% (3) | |
| Books | 2% (2) | |
| Other | 3% (3) | |

| **Q.21: What content areas do you edit? Please select all that apply.** Selected choices. (*n*=119) | | |
|---|---|---|
| Biomedical science | | 85% (101) |
| Biology | | 57% (68) |
| Public health | | 57% (68) |
| Engineering | | 26% (31) |
| Chemistry | | 24% (29) |
| Humanities | | 24% (28) |
| Physics | | 18% (21) |
| Other (please specify) | | 28% (33) |

| **Q.22a: What percentage of your time do you spend on the following activities? (Should total 100% and equate to the total amount of time worked.)** Categorized responses – note that percentages shown below for each activity only represent the percentage of respondents who indicated any portion of time spent on that activity, with further variation in what fractions of time were spent on each activity, as shown for Editing in Q.22b. (*n*=112) | | |
|---|---|---|
| Editing (e.g., editing, related correspondence, record keeping) (see Q.22b) | | 100% (112) |
| Administrative work (e.g., billing, reports) | | 73% (82) |
| Mentoring | | 46% (52) |
| Teaching | | 38% (43) |
| Communications (e.g., newsletter, blog, podcast) | | 36% (40) |
| Other (please specify) | | 35% (39) |
| Management/supervision | 25% (28) | |
| Grant coordination | 21% (23) | |
| Research assistance (e.g., systematic reviews) | 15% (17) | |
| Publication management (e.g., journal managing editor) | 11% (12) | |

| **Q.22b: Percent of time spent on editing (derived from data in Q22a).** (*n*=112) | |
|---|---|
| ≤10% | 1% (1) |
| 11-20% | 4% (5) |
| 21-30% | 8% (9) |
| 31-40% | 11% (12) |
| 41-50% | 21% (23) |

| | |
|---|---|
| 51-60% | 10% (11) |
| 61-70% | 8% (9) |
| 71-80% | 20% (22) |
| 81-90% | 8% (9) |
| 91-100% | 10% (11) |

| **Q.23: Approximately how many small (e.g., 1 page or less) editing projects do you work on in a year (including revisions of the same project)?** Categorized responses. (*n*=114) | |
|---|---|
| 0 | 20% (23) |
| 1-10 | 32% (36) |
| 11-20 | 13% (15) |
| 21-30 | 4% (5) |
| 31-40 | 4% (5) |
| 41-50 | 12% (14) |
| 51-60 | 2% (2) |
| 61-70 | 0% (0) |
| 71-80 | 1% (1) |
| 81-90 | 1% (1) |
| 91-100 | 1% (1) |
| >100 | 4% (5) |
| Other | 5% (6) |

| **Q.24: At your current employer, what career pathways or opportunities are available to you in your role? Please select all that apply.** Selected Choices. (*n*=111) | |
|---|---|
| Manager/Director Track | 34% (38) |
| Scientific Editor Track | 31% (34) |
| I am a freelancer | 26% (29) |
| Administrator Track | 13% (14) |
| Editorial Track (e.g., at a journal) | 6% (7) |
| Faculty Track | 6% (7) |
| Other | 4% (4) |
| Unsure | 14% (15) |
| None | 13% (14) |

| **Q.25: When did you begin your career in scientific editing?** Categorized responses. (*n*=127) | |
|---|---|
| <1 year ago | 2% (2) |
| 1-5 years ago | 10% (13) |
| 6-10 years ago | 20% (26) |
| 11-15 years ago | 25% (32) |
| 16-20 years ago | 15% (19) |
| >20 years ago | 28% (35) |

| **Q.26: At what stage in your education and working life did you begin your career as a scientific editor?** Selected choices. (*n*=126) | |
|---|---|
| During undergraduate school | 1% (1) |
| After undergraduate school | 5% (6) |
| After undergraduate school and work experience | 21% (26) |
| During graduate school | 3% (4) |
| After graduate school | 10% (13) |
| After graduate school and work experience | 27% (34) |
| During postdoctoral fellowship | 1% (1) |
| After a postdoctoral fellowship | 11% (14) |
| After a postdoctoral fellowship and work experience | 11% (14) |
| After a faculty position | 5% (6) |
| After other unrelated career | 6% (7) |

1

| **Q.27: How did you first hear about scientific editing as a profession?** Selected choices. (*n*=123) | | |
|---|---|---|
| Informal work for colleagues that expanded into a formal position | | 27% (33) |
| From a friend, colleague, or mentor | | 24% (30) |
| Intentional job search for related positions | | 22% (27) |
| Started as part of another job duty | | 6% (7) |
| Intentional editing job search | | 5% (6) |
| Recruited | | 4% (5) |
| Presentation on scientific editing or related topic | | 4% (5) |
| Ad (newspaper or social media or other) | | 4% (5) |
| Other (please specify) | | 4% (5) |
| Do not know | 1% (1) | |
| Job fair | 1% (1) | |
| Internship opportunity | 1% (1) | |
| Article about scientific editing | 1% (1) | |
| College advisor or career counselor | 1% (1) | |

2

| **Q.28: What professional organizations do you belong to or have you belonged to in the past 10 years? Please select all that apply.** Selected choices. (*n*=127) | |
|---|---|
| Board of Editors in the Life Sciences | 46% (59) |
| American Medical Writers Association | 39% (50) |
| National Organization of Research Development Professionals | 36% (46) |
| Scientific Editors Network | 35% (45) |
| Council of Science Editors | 23% (29) |
| Editorial Freelancers Association | 13% (16) |
| Copyediting-L | 6% (8) |
| National Association of Science Writers | 5% (6) |
| Northwest Editors Guild | 4% (5) |
| International Society for Medical Publication Professionals | 3% (4) |

| | |
|---|---|
| European Medical Writers Association | 3% (4) |
| Editors Tea Club | 2% (2) |
| Other | 4% (5) |

| **Q.29: When you were getting started in scientific editing, what resources were helpful to you? Please select all that apply.** Selected choices. (*n*=125) | |
|---|---|
| Colleagues or on-the-job training | 74% (93) |
| Written material | 59% (74) |
| Professional organizations | 46% (58) |
| Workshops or seminars | 42% (52) |
| Mentoring | 27% (34) |
| Grant administration | 12% (15) |
| Student activities | 2% (3) |
| Other | 10% (12) |

| **Q.30: What training and resources do you have access to that are supported by your employer? Please select all that apply.** Selected choices. (*n*=88, excluding 26 respondents categorized as primarily self-employed) | |
|---|---|
| Subscriptions to reference books, journals, or websites | 74% (65) |
| Access to written guidelines or style guides | 70% (62) |
| Software | 70% (62) |
| Paid membership in professional societies | 63% (55) |
| Internal presentations or workshops | 61% (54) |
| Funding to attend local or national conferences | 61% (54) |
| Other | 11% (10) |

| **Q.31: How did you acquire your first job in scientific editing? Please select all that apply.** Selected choices. (*n*=127) | |
|---|---|
| Through a posted job opportunity | 54% (68) |
| Started freelancing | 18% (23) |
| Through a friend or a colleague | 15% (19) |
| Started editing informally for colleagues and it was expanded into a formal position | 9% (11) |
| Was recruited | 7% (9) |
| Position evolved or was created within other employment situation | 6% (7) |
| Internship | 1% (1) |
| Applied for different job, but was hired as editor | 2% (3) |
| Promoted into position | 1% (1) |
| Cold approach to organization | 1% (1) |

| **Q.32: What resources or training (e.g., workshops, networking opportunities) do you wish had been available to you before becoming a scientific editor or early in your career?** Categorized responses. (*n*=109) | |
|---|---|
| Workshops and webinars | 23% (25) |

| | |
|---|---|
| Satisfied/NA | 22% (24) |
| Exposure, mentorship, and training | 19% (21) |
| Networking and conferences | 17% (18) |
| Coursework or certification | 12% (13) |
| Literature and style guides | 11% (12) |
| Professional organizations | 7% (8) |
| Journal access, software, and online tools | 0% (0) |
| Unsure | 2% (2) |
| Other | 8% (9) |

1

| **Q.33: What skills do you think are important for a successful career in scientific editing?** Categorized responses. (*n*=109) | |
|---|---|
| Efficient work method | 55% (60) |
| Interpersonal and client relations | 49% (53) |
| Language and communication skills | 46% (50) |
| Emotional intelligence | 35% (38) |
| Subject-matter expertise | 35% (38) |
| Critical thinking skills | 25% (27) |
| Familiarity with funding orgs/publishers | 13% (14) |
| Technical/clinical skills | 6% (6) |
| Business aptitude | 5% (5) |

2

| **Q.34: What resources for scientific editing do you wish were available to you now?** Categorized responses. (*n*=110) | |
|---|---|
| Satisfied/NA | 34% (37) |
| Literature and style guides | 16% (18) |
| Journal access, software, and online tools | 15% (16) |
| Networking and conferences | 15% (16) |
| Other | 15% (16) |
| Coursework or certification | 5% (5) |
| Professional organizations | 4% (4) |
| Unsure | 4% (4) |
| Exposure, mentorship, and training | 2% (2) |
| Workshops and webinars | 0% (0) |

3

| **Q.35: What is your favorite source for professional development or support**? Categorized responses. (*n*=102) | | |
|---|---|---|
| Professional organizations | | 76% (78) |
| National Organization of Research Development Professionals | 20% (20) | |
| Scientific Editors Network (ScENe) | 20% (20) | |
| American Medical Writers Association (AMWA) | 18% (18) | |
| Board of Editors in the Life Sciences (BELS) | 14% (14) | |
| Other/unspecified | 11% (11) | |

| | | |
|---|---|---|
| Local organizations | 6% (6) | |
| ACES: The Society for Editing | 4% (4) | |
| Editorial Freelancers Association (EFA) | 4% (4) | |
| Council of Science Editors (CSE) | 3% (3) | |
| Online resources | | 9% (9) |
| Colleagues | | 7% (7) |
| Office/internal resources | | 4% (4) |
| None | | 2% (2) |
| Style guides | | 1% (1) |
| Other | | 1% (1) |

| **Q.36: What resources do you use on a daily basis? Please select all that apply.** Selected choices. (*n*=113) | |
|---|---|
| Journal or funding agency websites | 67% (76) |
| Style guides | 61% (69) |
| Templates | 38% (43) |
| Books/workbooks | 22% (25) |
| Other | 19% (22) |

| **Q.37: Approximately how many clients do you work with in a year?** Categorized responses. (*n*=113) | |
|---|---|
| 1-10 | 20% (23) |
| 11-20 | 22% (25) |
| 21-30 | 19% (22) |
| 31-40 | 5% (6) |
| 41-50 | 7% (8) |
| 51-60 | 6% (7) |
| 61-70 | 1% (1) |
| 71-80 | 1% (1) |
| 81-90 | 0% (0) |
| 91-100 | 4% (4) |
| >100 | 4% (4) |
| Other | 11% (12) |

| **Q.38: Do you work for an individual unit (e.g., services a college or department) or central office (e.g., services the entire university).** Categorized responses. (*n*=63) | |
|---|---|
| Department within the university/institution | 37% (23) |
| Central office (service entire university) | 32% (20) |
| College within the university/institution | 17% (11) |
| Division within the university/institution | 14% (9) |

| **Q.39: Who do you report to?** Selected choices. (*n*=61) | |
|---|---|
| Head of department/institute | 36% (22) |

| | |
|---|---|
| Core director/associate director | 23% (14) |
| VP for research or similar | 18% (11) |
| Faculty member | 13% (8) |
| Dean of college | 10% (6) |

1

| **Q.40: What type of editing do you do? Please select all that apply.** Selected choices. (*n*=119) | | |
|---|---|---|
| Substantive editing for clarity and flow | | 97% (115) |
| Grammar/mechanics | | 93% (111) |
| Strategy and grantsmanship | | 58% (69) |
| Review of the science | | 45% (53) |
| Compliance | | 45% (54) |
| Other (please specify) | | 5% (6) |
| Styling/formatting | 3% (4) | |
| Fact-checking | 2% (2) | |

2

| **Q.41: Do you track outcomes for the projects you work on? If yes, please explain how.** Selected choices. (*n*=112) | | |
|---|---|---|
| Yes, I track outcomes | | 64% (72) |
| Track on own | 29% (32) | |
| Track informally | 12% (13) | |
| Surveys, inquiries, client feedback/follow up | 9% (10) | |
| Department tracks | 4% (5) | |
| Programmed alerts, check databases, review literature | 5% (6) | |
| No, I do not track outcomes | | 36% (40) |

3

| **Q.42: What is your method for determining the impact or success of your work? (e.g., funded proposals, accepted manuscripts, client testimonials, number of return clients, etc.).** Categorized responses. (*n*=105) | |
|---|---|
| Client satisfaction and retention | 71% (75) |
| Outcomes and rate of success | 58% (61) |
| Evaluation and review outcomes | 8% (8) |
| Manuscript and proposal submissions | 6% (6) |
| Service volume and activity metrics | 5% (5) |
| Internal evaluations and professional standards | 5% (5) |
| Sales, values, and training initiatives | 4% (4) |
| None | 5% (5) |

4

| **Q.43: If you currently run an office or business, how did you learn the business/administrative aspects needed for this position?** Categorized responses. (*n*=39) | |
|---|---|
| Self-taught | 49% (19) |
| Online | 23% (9) |
| Coursework | 21% (8) |
| Prior experience | 13% (5) |

| | |
|---|---|
| Consultation | 13% (5) |
| Job training | 10% (4) |
| Books | 8% (3) |
| Workshops | 5% (2) |
| Other/unspecified | 5% (2) |
| Social media | 3% (1) |
| Editorial Freelancers Association (EFA) resources | 3% (1) |

1

| **Q.44: What are your three favorite aspects of being a scientific editor?** Categorized responses. (*n*=110) | |
|---|---|
| Intellectually interesting | 67% (74) |
| Helping someone | 48% (53) |
| Working with smart/great people (faculty/editors) | 42% (46) |
| Work-life flexibility | 35% (38) |
| Meaningful contribution to science | 29% (32) |
| Suited to strengths | 21% (23) |
| Good compensation/steady work | 5% (5) |
| Clear endpoints | 2% (2) |

2

| **Q.45: What are your three least favorite aspects of being a scientific editor?** Categorized responses. (*n*=106) | |
|---|---|
| Clients | 48% (51) |
| Timing issues | 38% (40) |
| Certain aspects of editing projects | 36% (38) |
| Feeling undervalued/lack of advancement | 24% (25) |
| Administrative aspects | 16% (17) |
| Inherent qualities of performing editing | 11% (12) |
| Feeling isolated | 10% (11) |
| Inherent challenges with publishing/funding landscape | 7% (7) |
| Technical issues | 6% (6) |
| Unclear impact | 5% (5) |
| Problems at institution | 5% (5) |
| Other editors | 2% (2) |
| Other | 3% (3) |

3

| **Q.46: How many editing projects >1 page do you work on in a year (including revisions of the same project)?** Categorized responses. (*n*=113) | |
|---|---|
| 0 | 0% (0) |
| 1-10 | 12% (13) |
| 11-20 | 13% (15) |
| 21-30 | 11% (12) |
| 31-40 | 9% (10) |
| 41-50 | 14% (16) |

| | |
|---|---|
| 51-60 | 6% (7) |
| 61-70 | 2% (2) |
| 71-80 | 4% (4) |
| 81-90 | 0% (0) |
| 91-100 | 7% (8) |
| >100 | 17% (19) |
| Other | 6% (7) |

| **Q.47: Related to scientific editing, what previous titles have you held (e.g., Senior Editor, Grant Writer, Research Development Officer, Administrative Operations Manager, Freelance Editor).** Categorized responses. (*n*=96) | |
|---|---|
| Title includes: “Editor” | 69% (66) |
| Title includes: “Writer” | 26% (25) |
| Title includes: “Research Development” | 7% (7) |
| Title includes: “Grant” | 20% (19) |
| Title includes: “Research” (but not “Development”) | 17% (16) |
| Title includes: “Science” or Scientific” | 31% (30) |
